# Weak pre-before-post pair biases sort synaptic weights in a conductance-based model of spontaneous network bursting

**DOI:** 10.64898/2026.09.03.749248

**Authors:** Filippo Groppi, Jessica Ferrari

**Affiliations:** Luviner, Parma, Italy; Università di Parma, Parma, Italy

**Author notes:** Corresponding author: Filippo Groppi, (Luviner).

**Keywords:** synaptic plasticity, STDP, spontaneous network bursts, computational neuroscience, stomatogastric ganglion model, preregistration, multi-electrode arrays

## Abstract

Spontaneous population bursts repeatedly expose synapses to correlated spike activity, but it is unclear whether such events reinforce pre-existing synaptic weight differences when precise firing order carries little information. We asked this in a conductance-based model of a bursting neuronal culture: 48 excitatory and 12 inhibitory stomatogastric-ganglion (STG) model neurons with sparse recurrent connectivity, short-term depression, and pair-based spike-timing-dependent plasticity (STDP), with every prediction and falsifier committed to a version-controlled record before the corresponding run. Across two cohorts of 10 connectivity wirings — realizations of one model, not independent preparations — stronger synapses consistently received greater net potentiation than weaker ones, although the preregistered magnitude criterion was not met. The spike statistic most closely associated with this sorting was not first-spike latency: stronger synapses instead showed a very small but highly consistent excess of pre-before-post spike pairs, 0.2–0.35 percentage points, accumulated over ∼10⁴ pairs per synapse in 60 s (median 8,960 in the out-of-sample cohort; ∼3 × 10⁶ per wiring). In a preregistered intervention, shortening bursts by 41% at approximately matched burst rate increased the sorting measure by about an order of magnitude in all five wirings. A second, physiologically distinct perturbation reproduced the relationship with realized burst duration, but duration could not be separated from per-burst spike count. All interventional evidence is computational. We then preregistered an observational test of the corresponding prediction in 16 archived recordings of developing cortical cultures. The predicted strength-ordered pair excess was not detected, and the predicted negative relationship with burst duration was absent. The former is a bounded null — the recordings exclude effects above +0.32 percentage points but cannot resolve the model’s ∼0.2-point magnitude; the latter was adequately powered under the preregistered criterion. A consistent positive latency–strength relationship, unpredicted by the model, was observed. The study therefore establishes a reproducible sorting phenomenon within a specific computational model and a testable burst-shortening prediction, but does not establish that the mechanism operates in living cortical networks; an intervention on cultured networks with an independently validated measure of synaptic strength is the decisive test.

## 1. Introduction

### 1.1 Question

Spontaneous, synchronized population bursts dominate the activity of developing and cultured neuronal networks (Wagenaar et al., 2006). What, concretely, does a burst do to an existing distribution of synaptic weights? The question is usually answered at the level of plasticity-rule algebra rather than measured in a preparation where the weight order is known and every spike is observable. Whether a related process contributes to burst-rich activity in vivo is unknown, and nothing in this study speaks to it directly.

### 1.2 Competing mechanisms

A natural hypothesis is temporal: strongly driven cells should fire earlier within a burst, and STDP should convert that order into selective potentiation. A competing hypothesis is statistical: bursts are noisy enough that any single event carries almost no signal, and sorting, if it happens, is an accumulation across very many weakly biased events. These hypotheses make different predictions about where the signal lives — in first-spike latencies, or in pair-ordering counts — and about what should modulate it.

Our question is adjacent to, but distinct from, recent burst-dependent plasticity theory (Naud & Sprekeler, 2018; Payeur et al., 2021), which treats bursts as a multiplexed code for credit assignment; here we ask the converse — not what bursts signal, but what spontaneous bursts do to an existing weight distribution. “Counting, not timing” is not new as abstract theory: rate-based reductions of STDP (Kempter, Gerstner & van Hemmen, 1999), its mapping onto BCM-type rules (Izhikevich & Desai, 2003; Bienenstock, Cooper & Munro, 1982) and calcium-based accumulation models (Graupner & Brunel, 2012) all describe plasticity as an integral over many weakly informative events. What is new here is showing it concretely in a bursting network with known weights, deriving a counter-intuitive intervention from it, and testing the observational consequence on real MEA recordings.

### 1.3 Why this model

We use crustacean STG model neurons not as a stand-in for cortical physiology but for two reasons. First, transparency: the Prinz et al. (2003) database is a validated, standardized parameter space, and adopting it removes the hand-tuned degrees of freedom that make small-network simulation results hard to trust. Second, the claim under test is algorithmic rather than physiological: pair-based STDP provides a tractable test case for whether weak spike-pair biases can accumulate during spontaneous bursts, and the hypothesis is that, under pair-based STDP, weak statistical biases in spike-pair ordering can accumulate into weight sorting even when event-level firing order carries little information. One structural difference is stated at the outset: bursting in this model is intrinsically paced by the cells’ own conductances, whereas bursts in cortical cultures are recurrence-driven (Section 4). Section 5 is the testable hypothesis that the principle transfers to cortical preparations — not the assertion that it does.

### 1.4 Study design and preregistration

We tested both hypotheses in this simulated culture under a preregistration discipline unusual for simulation work: predictions, priors, and falsifiers were committed to a version-controlled laboratory record before each simulation ran, and negative or partial outcomes are reported as they were scored rather than reframed (Methods, Section 7.4; deviations and corrections in Section 6).

The temporal hypothesis was not supported under the preregistered criterion. The statistical hypothesis received directional support in out-of-sample simulation and in two computational interventions, and acquired an unexpected refinement: shorter bursts sort more strongly. We then subjected the prediction to its own test: its observational half, committed before download, was tested on the public Wagenaar et al. (2006) recordings and was not reproduced within the stated bounds (Section 2.6) — which recasts the interventional half as the decisive experiment.

## 2. Results

We reserve *replication* for the out-of-sample cohort; within a cohort, wirings are independent connectivity realizations of one model specification, not independent preparations. Two claims were registered jointly on ten wirings, hierarchically bound so that the mechanism claim could not rescue the outcome claim. The direction held in all ten realizations of each cohort (Sections 2.2 and 2.3); the magnitude criterion was not met (effect/spread = 0.92) and is reported as failed.

### 2.1 Burst-scale first-spike latency does not encode synaptic strength

If strongly driven cells fire earlier, Spearman ρ between a cell’s median within-burst first-spike latency and its total incoming weight should be negative in ≥8/10 wirings. Result: negative in 5 of 10, mean ρ = −0.014 ± 0.176 (Figure 2A). The preregistered criterion was not met; the temporal hypothesis was not supported under the preregistered criterion. We found no evidence that burst-scale first-spike latency tracked synaptic weight in this preparation.

**Figure 1.**
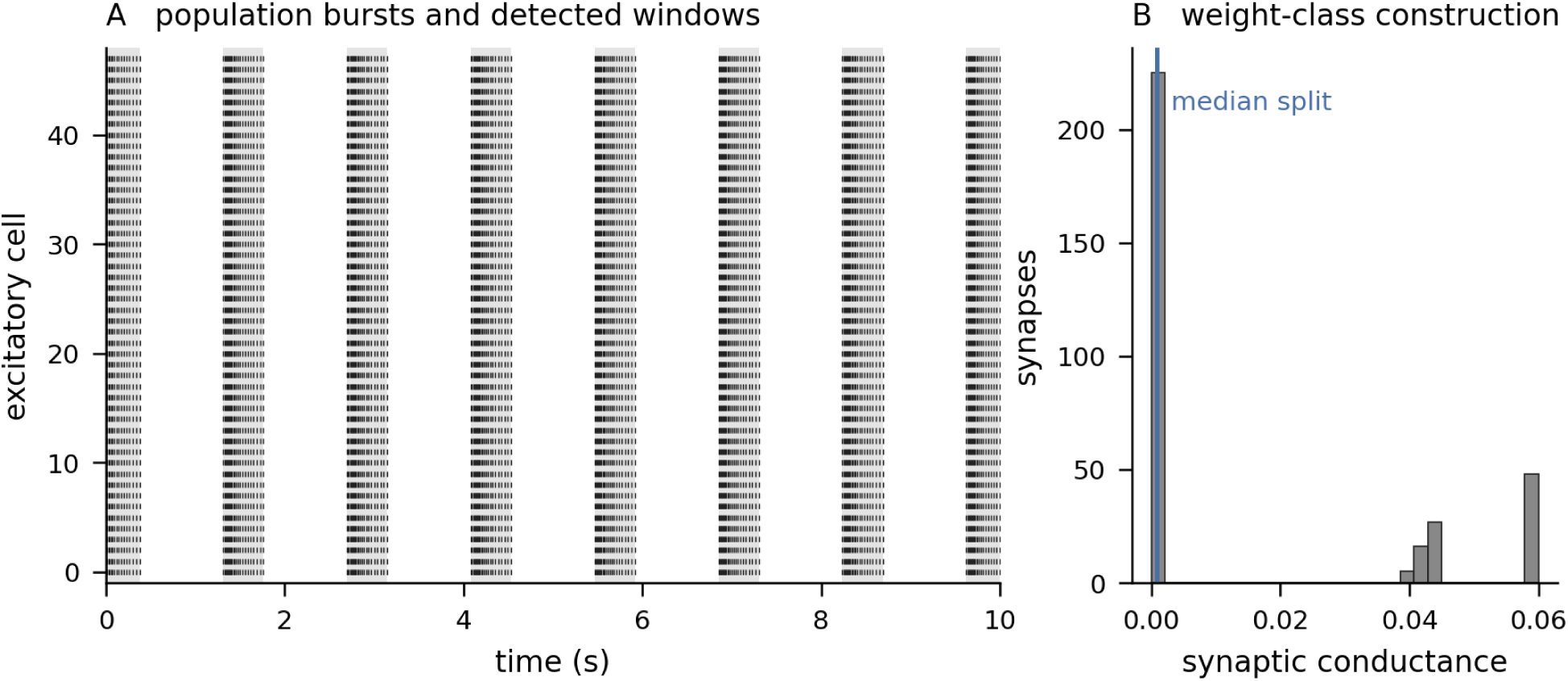
The preparation and its instruments. (`fig1_preparation.pdf`, from `stg_pm_s*.json`.) **A**, raster of the 48 excitatory cells over 10 s of a 60 s run, with the detected burst windows shaded; bursts are delimited at full width at 20% of each burst’s own peak. **B**, the distribution of plastic recurrent conductances at the end of the same run, with the median split that defines the strong and weak classes. A wiring in the modal bursting regime is shown; the two off-regime wirings identified in Section 2.2 differ in burst count and in the fraction of weights driven to zero, and would not be representative.

**Figure 2.**
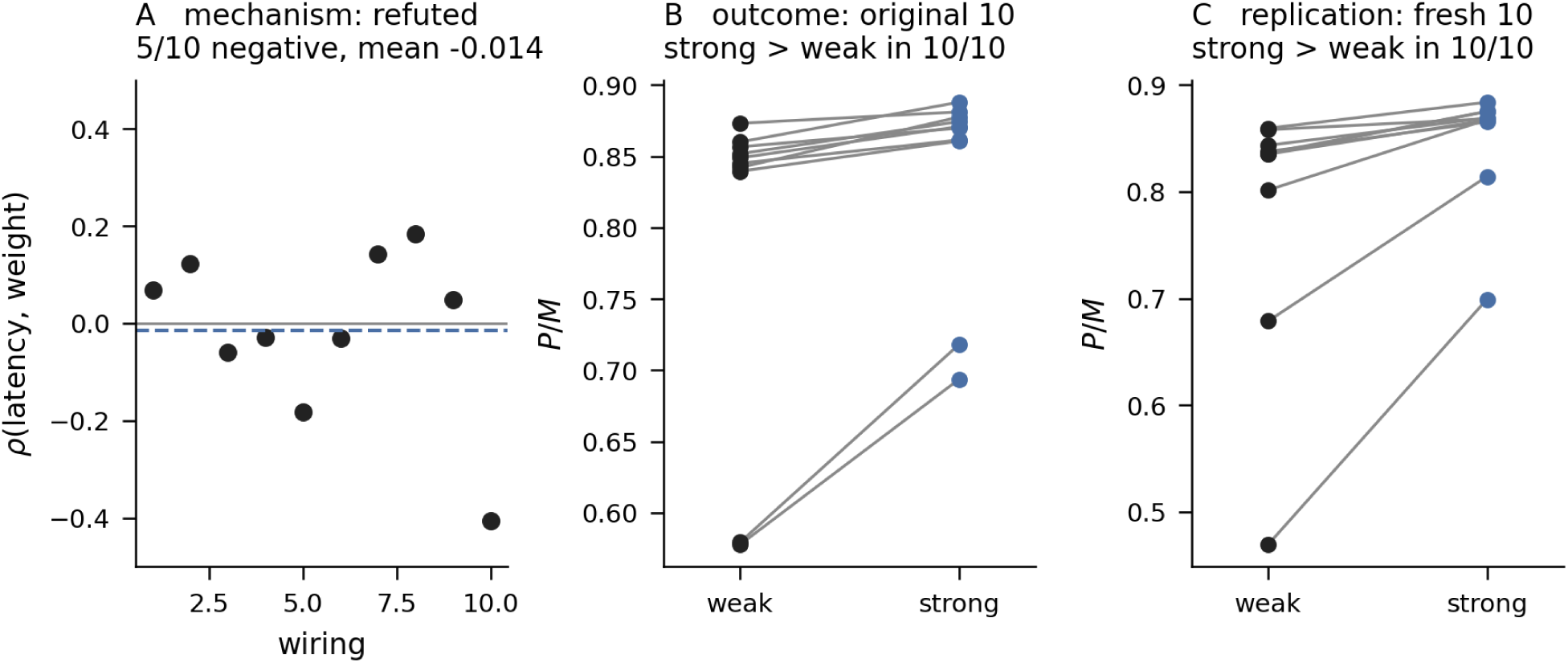
The temporal mechanism is not supported; the directional outcome is. (`fig2_mechanism_outcome.pdf`, from `stg_pm_s*.json` and `stg_pm4_s*.json`.) **A**, Spearman ρ between a cell’s median within-burst first-spike latency and its total incoming weight, one point per wiring, against the requirement of ≥8/10 negative: 5 of 10 are negative and the mean (dashed) is −0.014. **B**, paired weak→strong P/M within each of the ten original wirings. **C**, the same on the ten wirings generated after the prediction was frozen. The magnitude criterion in **B** was not met and is reported as failed in Section 2.2; the direction is 10/10 in both cohorts.

### 2.2 Stronger synapses accumulate a small pre-before-post pair excess

#### Outcome: direction met in 10/10; the magnitude bar was not met

The strong-minus-weak P/M difference (Methods, Section 7.3) was positive in 10 of 10 wirings (exact two-sided binomial p = 0.00195; this tests directional consistency across model realizations, not independent biological preparations; mean +0.0421, t = +2.90 as a secondary description; Figure 2B), but the criterion also required the effect to exceed the between-wiring spread, and it did not (effect/spread = 0.92). We report this as scored: the direction is consistent across all ten wirings; the magnitude varies across wirings by slightly more than its own mean.

#### The associated statistic, identified by pre-authorized re-analyses

Neither spikes per burst (strong 20.33 vs weak 20.50, same sign in 2/10) nor burst participation (0.894 vs 0.901, 2/10) differs by class. What differs, with the same sign in 10 of 10 wirings, is the pre-before-post pair fraction: 0.5147 (strong) vs 0.5125 (weak) — a 0.22-percentage-point excess of pre-before-post pairs on strong synapses, expressed over ∼10⁴ pre–post pairs per synapse in 60 s (median 8,960 per synapse in the out-of-sample cohort, 8,837–9,238 per wiring excluding the two off-regime wirings; ∼3 × 10⁶ pairs per wiring over ∼340 synapses) and amplified by the exponential STDP kernels into a P/M difference of a few percent. This also explains a previous null in the same instrument: an earlier preparation with 18–46 ms, one-to-two-spike events showed no sorting (+0.0041 ± 0.0432) — its events were not too brief to *have* an order, but too brief to *accumulate* one. Per-class fractions were not persisted per wiring for this original cohort; the out-of-sample replication of the statistic, which was, is shown in Figure 3A.

**Figure 3.**
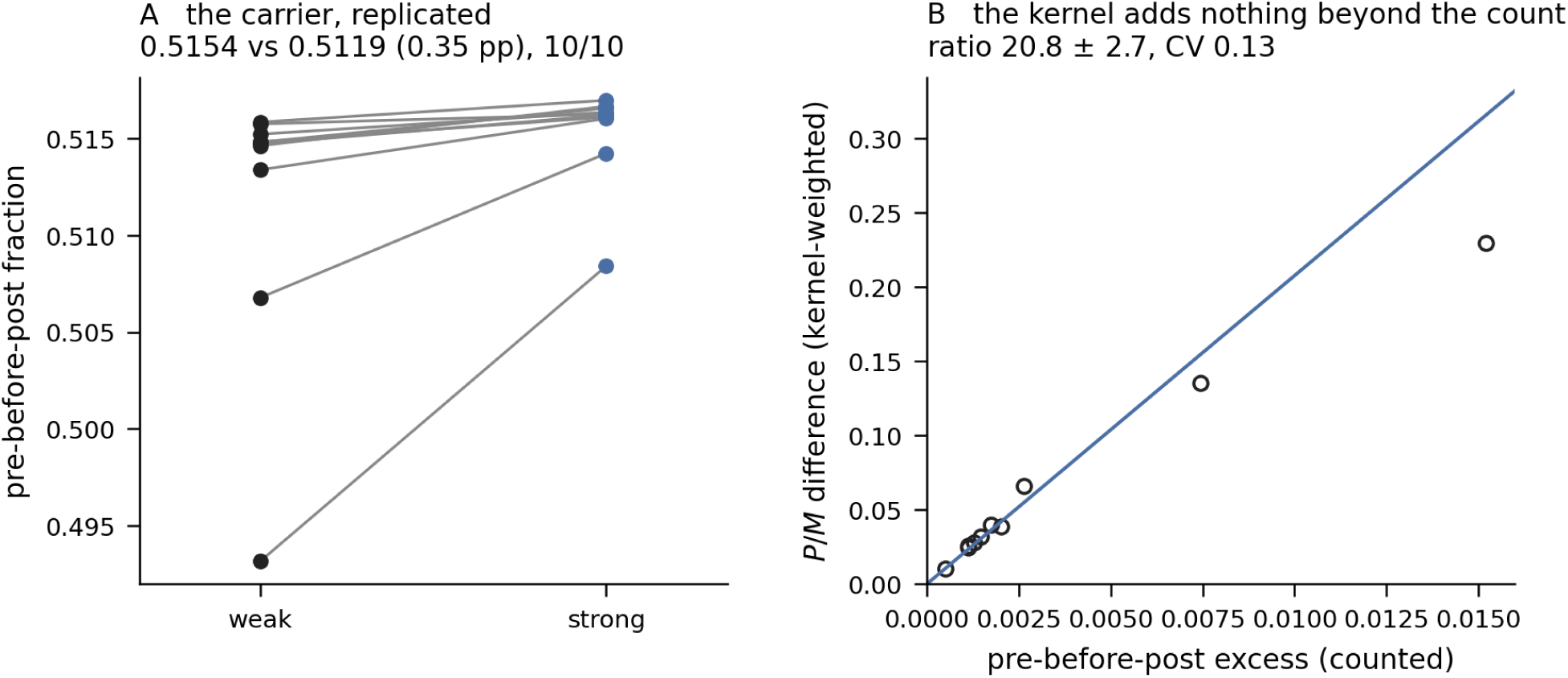
The associated statistic, replicated out of sample. (`fig3_carrier.pdf`, from `stg_pm4_s*.json`.) **A**, pre-before-post pair fraction by class in each of the ten fresh wirings: 0.5154 against 0.5119, a 0.35-percentage-point excess on strong synapses, same sign in 10 of 10. Per-class values were not persisted for the original cohort, so this figure shows the replication rather than the original result; the two cohorts’ excesses (0.22 percentage points in the original cohort, 0.35 in the replication) are independent estimates on different wirings. **B**, the kernel-weighted P/M difference against the counted pre-before-post excess, one point per wiring, with a proportional line through the origin at the mean per-wiring ratio (not a least-squares fit): the ratio is 20.8 ± 2.7 (CV 0.13), so the exponential STDP weighting contributes little beyond the count under the present rule.

### 2.3 The effect replicates in out-of-sample wirings

The direction was then registered as a prediction on ten wirings that did not exist when the prediction was written (registration pushed before the first simulation step). Result: the strong-minus-weak P/M difference and the pre-before-post excess were positive in 10 of 10 fresh wirings (P/M differences +0.010 to +0.230; exact two-sided binomial p = 0.00195, again a test of directional consistency across model realizations rather than of independent preparations; Figure 2C). The associated statistic replicated with them: the pre-before-post pair fraction was 0.5154 (strong) vs 0.5119 (weak), a 0.35-percentage-point excess, same sign in 10 of 10 (Figure 3A). The original and replication cohorts are different wirings, so the two excesses (0.22 percentage points in the original cohort, 0.35 in the replication) are two independent estimates of the same quantity rather than a discrepancy.

The correlation between the two measures across wirings (ρ = +0.988) is close to arithmetic, since both are computed from the same spike pairs — their ratio is constant to a coefficient of variation of 0.13 (Figure 3B). The replication is therefore **one** out-of-sample confirmation, not three. What the near-constant ratio genuinely establishes is that the effect is associated with **how many** pre-before-post pairs there are rather than with their fine timing within the STDP kernel: the exponential weighting contributes little beyond the pair count under the present STDP rule and activity regime. This claim concerns pair timing versus pair count under a pair-based rule; it does not speak to higher-order spike statistics, which triplet-STDP models were introduced precisely to capture (Pfister & Gerstner, 2006).

A post-hoc observation, explicitly not scored: the two wirings that deviated from the modal bursting regime had the largest sorting effects, and sorting magnitude correlated more strongly with burst duration (ρ = −0.47) than with burst count (+0.24) — suggesting, on n = 2, that shorter bursts sort more strongly. This anecdote was frozen as the *predictor* of the next experiment before that experiment ran.

#### 2.3.1 Exploratory, post-hoc checks [exploratory, post-hoc — not registered]

Three checks added after the results were seen, none in the PM-4 registration, all recomputed from persisted data with no new simulation (full text, Supplementary Results S3; Figures S1, S2). With weight as a continuous predictor, the rank correlation between a synapse’s weight and its pre-before-post fraction is ρ = +0.386, positive in 10 of 10 realizations, and the median curve rises with weight but is not monotone [exploratory]. A kernel ablation reproduces the across-wiring ratio CV at 0.132 and gives a per-synapse ρ(count fraction, P/M) = +0.977; the departure from proportionality is visible but not established [exploratory]. Applying the tissue test’s per-synapse burst-shuffle null to the model cohort gives a median excess over the null of +0.022 percentage points, positive in 8 of 10, sign test p = 0.109 [exploratory] — holding Sections 2.1–2.5 to the standard of Section 2.6 weakens them from 10 of 10 to 8 of 10, and we report this rather than omit it.

### 2.4 Shortening bursts increases sorting by about an order of magnitude

We fixed the direction in advance — shorter bursts sort *more strongly*, against the naive “more time to accumulate” reading — and manipulated burst duration synaptically, by raising inhibitory conductance g_ie (0.03/0.06/0.10) in five wirings, with a guard that burst-rate spread stay below 10% (measured: 8.6%, so rate is matched by construction; a failed guard would have voided that wiring × level cell only). Raising g_ie from 0.03 to 0.10 shortened bursts from 461 to 272 ms (−41%) and strengthened sorting in 5 of 5 wirings on both measures: the pre-before-post excess rose from +0.00095 to +0.00958 and the P/M difference from +0.02118 to +0.13209 — approximately an order of magnitude from a small baseline (10.1× and 6.2× respectively; the absolute values are the informative quantity). Three of five wirings were monotone across all three levels. A registered observational arm (natural duration variation at fixed g_ie) was uninformative rather than failed: surviving wirings at the middle level spanned 444–449 ms — a 1.0% spread — leaving nothing to correlate, which is precisely why an intervention was needed.

This experiment alone cannot attribute the effect to duration: g_ie is a confounded lever (there is no knob in this preparation that moves burst duration and nothing else), a limitation named in its registration.

### 2.5 A second perturbation: sorting increases as burst duration decreases, with burst duration and per-burst spike count remaining unresolved candidate mediators

To break the confound we registered a second intervention with disjoint physiology: lowering the slow calcium conductance g_CaS inside the excitatory cells (×0.8, ×0.6), touching no synapse, at fixed g_ie = 0.03. Predictions: sorting stronger at CaS ×0.6 than baseline within wirings; sorting tracking *realized* burst duration pooled across arms **and within each arm** (so the pooled correlation cannot be two clusters at different offsets); and a pre-specified ceiling on the claim (below).

Both arms share the same five wirings — the intrinsic lever was applied to the wirings of Section 2.4 — so this is a within-wiring triangulation using two physiologically distinct perturbations (same five wirings), not two independent tests on separate preparations.

The registered directional predictions received directional support in the surviving data, subject to the pre-specified exclusions and the unresolved duration/spike-count disjunction. CaS ×0.6 shortened bursts to 258–282 ms and strengthened sorting in 4 of 4 wirings with surviving data (4 of 5 overall; one wiring lost both intrinsic levels to network death, Section 7.6). Sorting tracked realized duration at ρ = −0.79 pooled (n = 18 de-duplicated cells; see Section 6 for the correction applied to the originally scored pool), −0.648 within the intrinsic arm, −0.808 within the inhibitory arm (Figure 4A). Figure 4 draws the shared baseline once. The within-arm correlations are unaffected by the correction.

**Figure 4.**
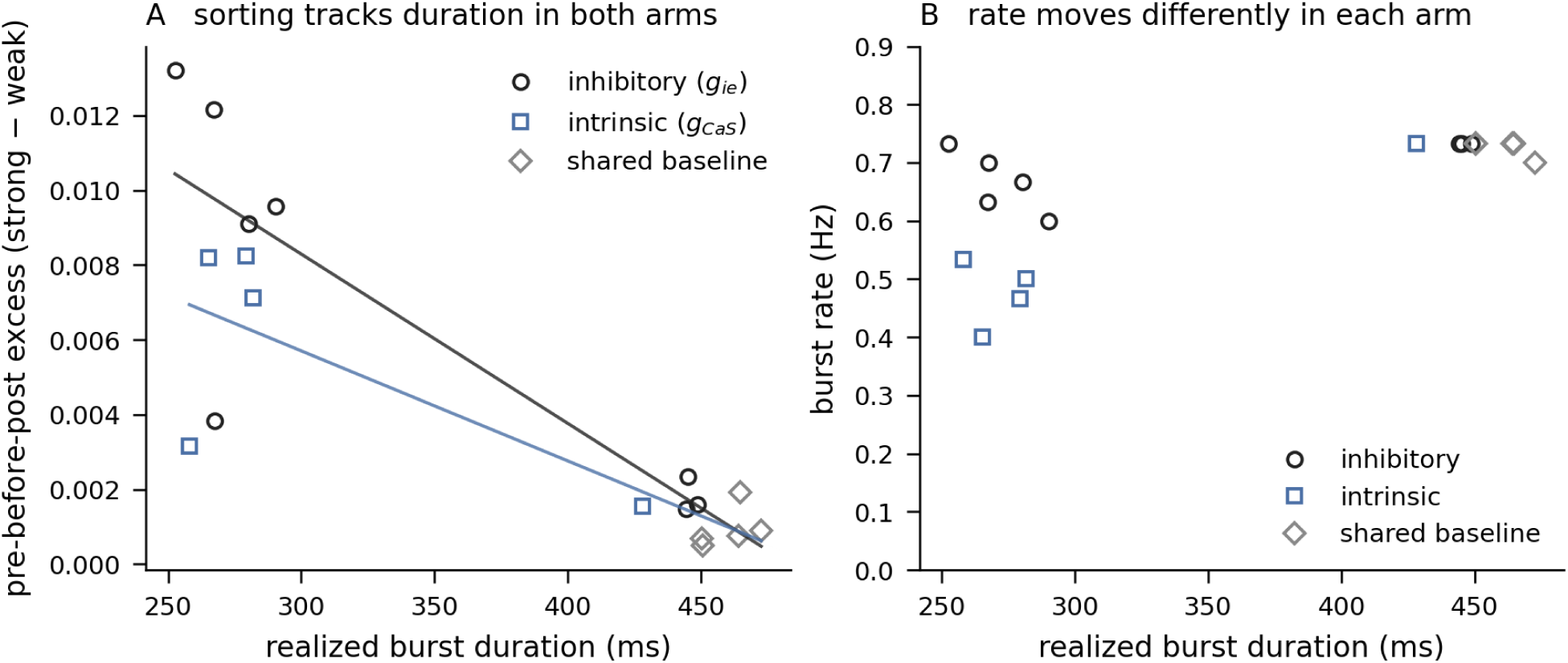
Two physiologically distinct perturbations on the same networks. (`fig4_interventions.pdf`, from `dur1_s*.json` and `dur2_s*.json`.) **A**, pre-before-post excess against realized burst duration for the inhibitory lever (g_ie, circles), the intrinsic lever (g_CaS, squares) and the baseline condition shared by both arms (diamonds), with a within-arm linear fit through each arm and its baseline. The shared points are the same runs — the intrinsic arm’s g_CaS ×1.0 *is* the inhibitory arm’s g_ie = 0.03 — and are drawn once, in their own marker, so that points are not double-counted. **B**, burst rate against realized duration for the same runs: rate is held nearly fixed across the inhibitory arm (8.6% spread) and falls substantially across the intrinsic arm, which is what excludes rate as the driver.

#### Rate is excluded as the driver by design rather than assumption

n the inhibitory arm, burst rate was held nearly fixed (8.6% spread) while sorting rose; in the intrinsic arm, rate *fell* substantially (0.73 → 0.40–0.53 bursts/s) while sorting rose the same way. No monotone function of burst rate can produce both arms’ trajectories (Figure 4B). This exclusion survives the shared-wiring caveat above, since it needs only the two rate profiles to disagree on the same networks.

#### The pre-specified ceiling stands

Spikes per burst falls with duration in both arms, and burst duration and spikes per burst were strongly collinear (ρ = +0.93 across the pooled cells), so the experiment cannot identify which variable mediates the effect; the registered tie-clause did not fire, but a bootstrap over the 18 cells shows the two candidates to be indistinguishable at this n (Supplementary Results S3). The strongest justified claim is, verbatim from the registration: the mediator is **“burst duration, or the spike count that falls with it.”** Separating them would require a lever that moves one without the other, absent in this model.

### 2.6 Test in archived cortical-culture recordings

We then tested the observational half of our own prediction on data we did not generate: the Wagenaar, Pine & Potter (2006) archive of spontaneous activity in developing dissociated cortical cultures on MEAs. Instruments, predictions, and falsifiers were committed before any file was downloaded.

#### Instruments

On an MEA there are no synaptic weights, and the standard strength estimator — a cross-correlogram peak — is essentially a pre-before-post pair count, so scoring “strong connections have more pre-before-post pairs” on the same spikes would be true by construction.

Strength (the rate-normalized cross-correlogram peak at |lag| ≤ 25 ms within bursts, median split) was therefore estimated on odd-numbered bursts and the excess scored on even-numbered ones, then swapped and averaged: the spikes that define a connection’s class are never the spikes that score it. The per-pair null is a burst shuffle (unit A’s spikes from burst *i* against unit B’s from burst *j*, 100 derangements), which preserves spike count, firing rate, and each unit’s typical position within a burst, and destroys only same-burst coordination; its conservative asymmetry is described in Supplement S1. The lag window was not inherited from the model: it is half of each recording’s median burst duration (92 ms on the probe recording), with burst duration measured by the dataset’s own published method, which our implementation reproduces (186 ms at 25 days in vitro against the published <200 ms). The scored perimeter is one culture from each of six plating batches at days in vitro {10, 17, 24} — batch siblings are not independent preparations, by the dataset authors’ own conclusion — with the first 300 s of each recording discarded and a guard voiding recordings with fewer than 8 bursts (two voided; N = 16; Section 7.6). All verdicts below are from the corrected scoring run (Section 6).

#### Verdicts, as scored

P1 (the strong half of putative connections shows a higher pre-before-post fraction in ≥ N−1 of N recordings): 6 of 16 (Figure 5A). P1 was not met. However, the preregistered threshold was substantially larger than the effect predicted by the model itself. Post-hoc power analysis showed that the dataset could exclude effects larger than +0.32 percentage points (mean −0.105 points, sd 0.80, 95% CI [−0.53, +0.32]) but could not distinguish absence of the mechanism from an effect of the approximately 0.2-point magnitude observed in the model; P1’s pass rule (15 of 16 positive) required a per-recording effect near 1.0 percentage point against a between-recording sd of 0.80. We therefore treat P1 as a bounded null rather than as evidence that the mechanism is absent. This mismatch between the preregistered decision threshold and the model-predicted effect was a registration error and is reported explicitly (Section 6). The null itself is calibrated rather than inflated: the median |z| of a recording against its own burst-shuffle null is 1.0 [exploratory], and the three recordings beyond 3σ of their null (+7.4σ, +6.0σ, −3.7σ) leave it in both directions.

**Figure 5.**
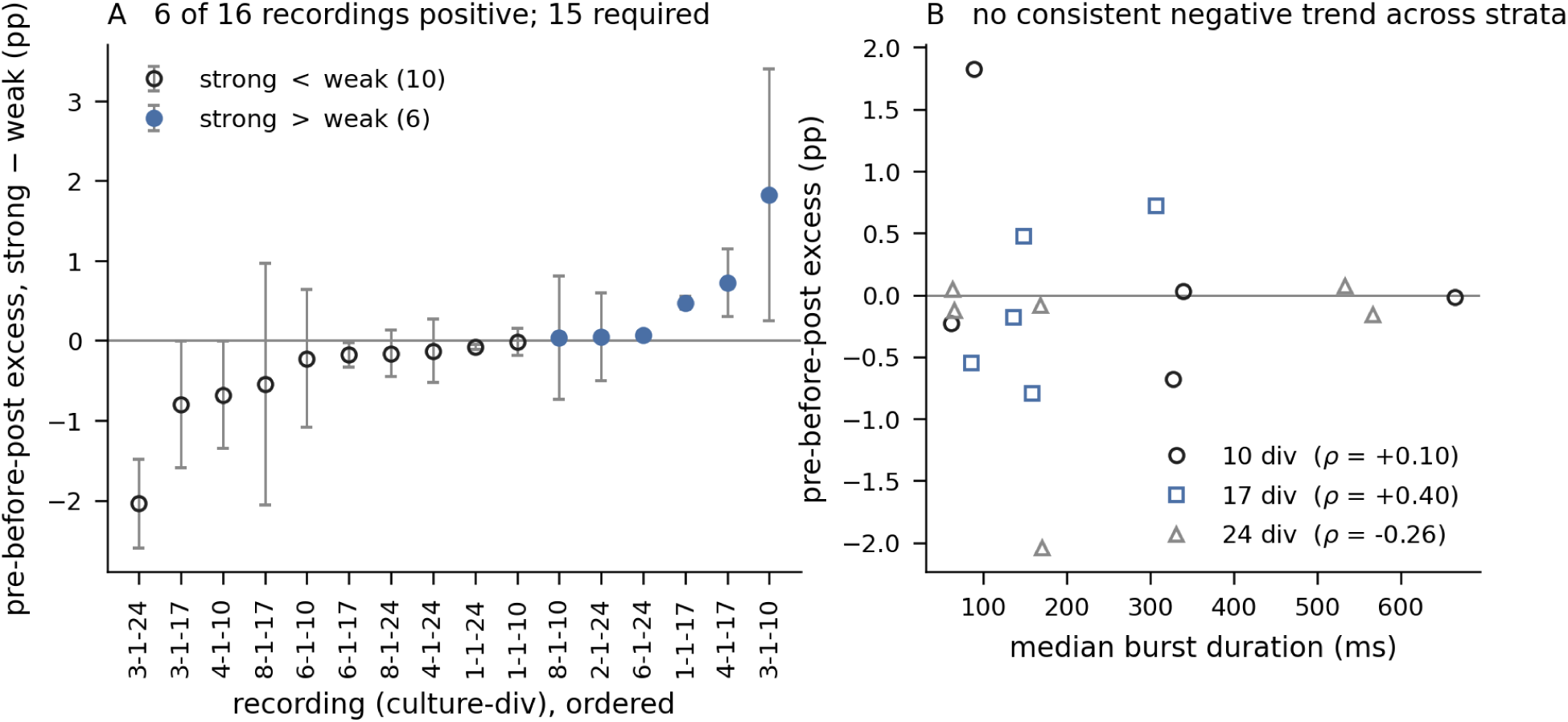
Test of the computational prediction in archived cortical-culture recordings. (`fig5_mea.pdf`, from the per-recording MEA-1 outputs; Wagenaar et al., 2006 data.) **A**, strong-minus-weak pre-before-post excess per recording against its own burst-shuffle null, one point per recording with the null spread: 6 of 16 positive against a bar of ≥15, mean −0.105 percentage points, 95% CI [−0.53, +0.32]. The model’s ∼0.2-point predicted excess lies inside the interval — the result is a bounded null. The null is calibrated rather than inflated: the median |z| of a recording against its own burst-shuffle null is 1.0, and the three recordings that leave their null by more than 3σ (6-1-24 at +7.4σ, 1-1-17 at +6.0σ, 3-1-24 at −3.7σ) leave it in both directions — what happens in tissue carries no consistent sign. **B**, sorting strength against median burst duration within age strata (days in vitro 10, 17, 24): no consistent negative trend across strata (ρ = +0.10, +0.40, −0.26 by stratum); the registered within-stratum test was not met on the pooled-within estimate (+0.08), with power.

**Figure 6.**
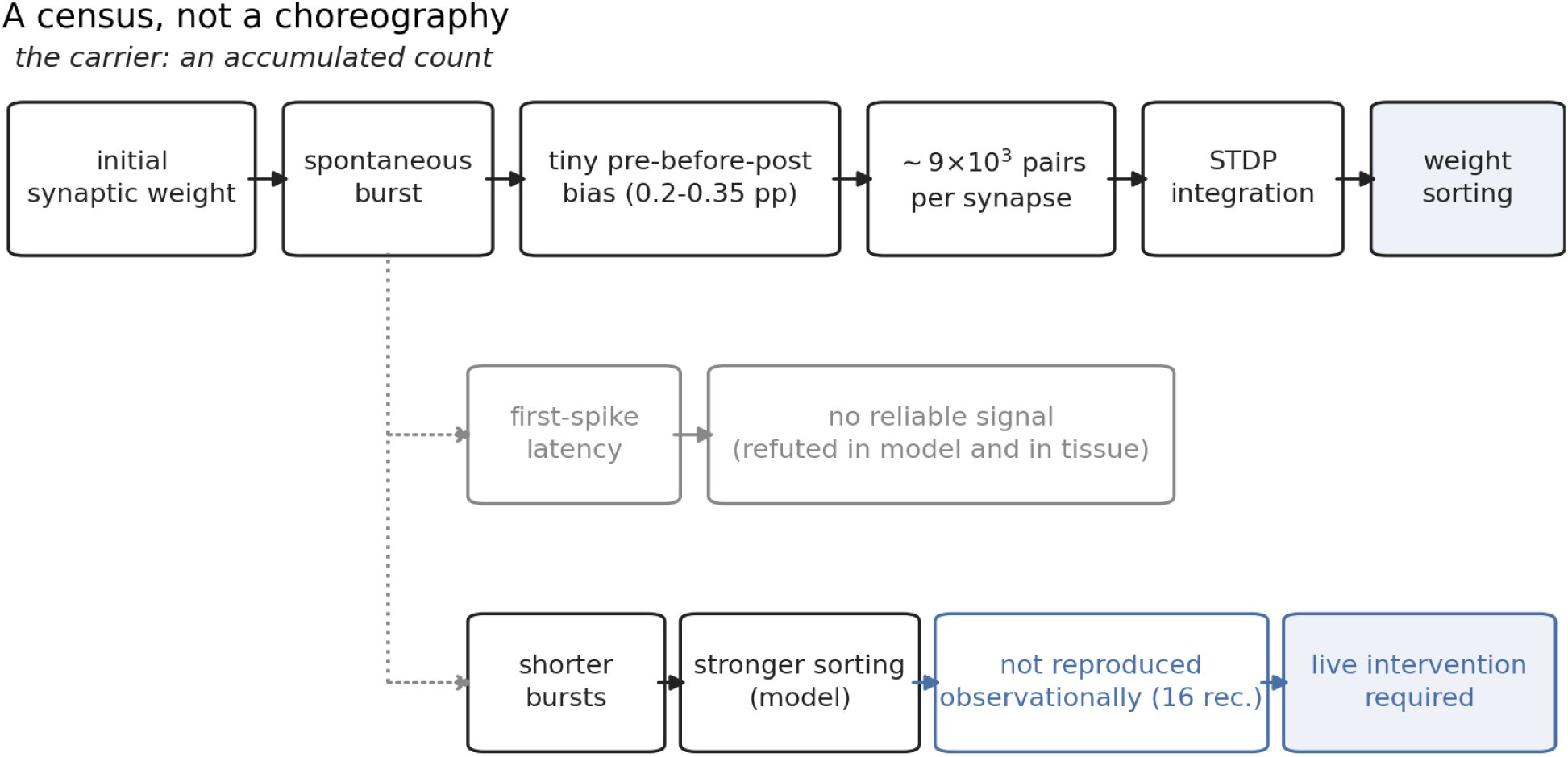
Model schematic. (`fig6_schematic.pdf`.) Three branches. *Main branch:* initial synaptic weight → spontaneous burst → tiny pre-before-post bias on stronger synapses (0.2–0.35 percentage points) → ∼10⁴ pairs per synapse in 60 s (∼3 × 10⁶ per wiring) → STDP integration → weight sorting. *Latency branch:* first-spike latency → no reliable signal (Section 2.1). *Intervention branch:* shorter burst → stronger sorting in the model (Sections 2.4–2.5) → not reproduced observationally in archived cortical recordings (Section 2.6) → live intervention required (Section 5). The model suggests a census rather than a choreography.

P2 (sorting strength correlates negatively with burst duration within age strata): ρ = +0.08 within strata (per-stratum +0.10, +0.40, −0.26; Figure 5B), not met on the merits — neither pre-specified escape applied: the underpower clause did not fire (within-stratum duration spread was 2.32× the across-stratum spread, nine times the floor), and the tie-clause did not fire (sorting tracked unit count at ρ = +0.05, *less* well than duration). P2 carries no bounded-null caveat: it had power by its own registered test and was not met on the merits. The asymmetry between P1 and P2 is a matter of what was committed: P2’s underpower criterion (within-stratum spread below 25% of across-stratum) was registered before download and did not fire, whereas no power criterion was registered for P1 and its minimum detectable effect was computed only afterwards.

P3 (first-spike-latency order carries no consistently negative signal): the registered criterion was met. But a consistent positive latency–strength relationship was found in tissue (ρ = +0.35, positive in 13 of 14 recordings [exploratory]), unexplained by this model and not scored under the registered, negative-only hypothesis; meeting the criterion is not evidence of latency irrelevance, and since P3 controlled nothing for firing rate, the positive relation cannot be attributed by this design.

## 3. Discussion

### Main computational finding

In this model, the weight order is read and reinforced by spontaneous bursts through a sub-percent excess of pre-before-post spike pairs, accumulated across hundreds of thousands of pairs and amplified by the plasticity kernel — while the intuitive temporal mechanism (strong cells fire first) received no support — across ten connectivity draws of a single model system and ten more, not independent preparations. Under the present STDP rule and activity regime the kernel’s fine timing contributes little beyond the pair count (post-hoc kernel ablation, Figure S2 [exploratory]); whether that holds under other kernels, time constants, or higher-order rules is untested. The model suggests a census rather than a choreography: sorting can emerge from a large number of weakly biased spike-pair events without a reliable burst-scale firing sequence. Whether a related mechanism contributes to burst-rich activity in vivo is unknown.

| Level of evidence | Result |
| --- | --- |
| Model | Sorting directionally reproducible (10 of 10 in each of two cohorts of connectivity realizations); magnitude criterion not met |
| Intervention in the model | Burst shortening amplifies sorting (~10×, 5 of 5); mediator (duration vs. spikes per burst) unresolved |
| Biological data | Observational prediction not reproduced: P1 a bounded null, P2 not met with power; an unpredicted positive latency–strength relation observed |

The direction remains informative despite the failed magnitude criterion because it held in all ten realizations of two independent cohorts and moved in the registered direction under intervention.

### Burst shortening

Shorter bursts sort more strongly — in the model. The direction was fixed before the data and is the opposite of the accumulation-time intuition; in tissue the corresponding observational relationship was not found — P2 was not met, with power, on the Wagenaar recordings (Section 2.6) — so this claim is, at present, a property of the model. A candidate account — long bursts saturate into undifferentiated firing whose late pairs dilute the informative early structure — is consistent with the spike-count collinearity but was not registered and is not claimed here. Burst duration and per-burst spike count remain unresolved candidate mediators: separating them would require a lever that moves one without the other, absent in this model; ρ(duration, spikes per burst) = +0.93 across the pooled cells, so no test at this n can separate them [exploratory].

### Biological test

The observational half of our prediction was not reproduced on real recordings (Section 2.6), and the two failures should be weighed differently. The duration relationship (P2) was not met with power and without escape clauses; on this dataset it is simply not there. The counting excess (P1) is a bounded null: the test could not resolve an effect at the model’s own scale, so absence and smallness remain unseparated. Three readings are compatible with the data. The mechanism may not transfer — the structural difference named in Limitations (intrinsically paced versus recurrence-driven bursting) is the candidate explanation, and it was stated before the test. The effect may exist at a scale this dataset cannot resolve, on electrode populations rather than synapses, under a null that deliberately reports a lower bound. Or the model may simply be wrong about tissue. These are not separable observationally: an intervention that shortens bursts at matched rate (Section 5) makes distinct predictions under each reading, which is why the interventional half is now the decisive experiment rather than a follow-up. The latency hypothesis fared differently in tissue than the model would suggest: no consistent negative relation appeared, which met the registered criterion, but a consistent positive latency–strength relationship did (ρ = +0.35, 13 of 14 [exploratory]), which this model does not explain and which was not scored.

### What is actually established

A reproducible, direction-consistent sorting phenomenon within one model specification, associated with a small pre-before-post pair bias rather than with firing order, and amplified by burst shortening under two physiologically distinct perturbations — together with a preregistered prediction whose observational form was not reproduced in archived cortical recordings. Not established: that the mechanism operates in living cortical networks, that duration rather than spike count is the mediator, or that the result is robust to the plasticity rule and the fixed potentiation/depression balance (Section 4). Every scored claim in this paper had its criterion, direction, and falsifier committed to version control before the number existed; failed clauses (the mechanism prediction, the magnitude bar in Section 2.2, and both observational predictions in Section 2.6) are reported as failures.

## 4. Limitations

All positive results are from simulation; the one test on real recordings (Section 2.6) is observational, and the predicted effect was not reproduced within stated bounds. The neurons are crustacean STG models, not cortical cells, and the network is small (48+12). A structural difference between the preparations must be stated plainly: cortical culture bursts are recurrence-driven, while bursting here is intrinsically paced — the excitatory conductance set is an endogenous burster, an intrinsic conductance controls burst duration (Section 2.5), and the recurrent synapses do not recruit activity at rest. Whether pair-counting sorting survives the move from intrinsically paced to recurrence-driven bursting is exactly what the Section 5 experiment would decide; the observational result of Section 2.6 is consistent with it not surviving, but cannot establish that. The tissue test carries its own limits, stated in its registration: a Wagenaar-era “unit” is an electrode — multi-unit activity from a population within ∼200 µm, not a neuron — so its P1 tested ordering between electrode populations, where a correlogram peak can reflect shared drive rather than a synapse; and its null deliberately subtracts static phase structure, so the measured excess is a lower bound (Supplement S1).

Plasticity is pair-based STDP with fixed kernels, and the rule itself may be the limit: triplet or voltage-based rules (Pfister & Gerstner, 2006) weight higher-order spike statistics that a pair rule cannot see, and could alter the result. The fixed potentiation/depression balance (the STDP window integral, Methods Section 7.2) is a modelling decision whose sensitivity was not tested. Because the fixed LTP/LTD balance was not varied, the sensitivity of the observed sorting to this parameter remains unknown, and no robustness is claimed. The construction is metaplasticity-adjacent (Abraham, 2008): a fixed LTP/LTD balance acts as the threshold against which activity is read. We likewise model no homeostatic scaling (Turrigiano & Nelson, 2004), the mechanism most likely to interact with — or oppose — the sorting described here over longer timescales. Real cortical and hippocampal neurons also carry calcium-dependent burst generation and dendritic spikes that a single-compartment model omits. Runs last 60 s of biological time, so the results describe the initial phase of short-timescale sorting, not a steady state — slower processes are invisible by construction.

Duration and per-burst spike count are collinear in every arm run so far; the mediator claim is bounded accordingly (Section 2.5). The magnitude criterion in Section 2.2 failed as pooled and stands as failed. A split noticed after the fact — not registered, therefore not scored — is informative about why: wirings in the 17-burst regime show +0.1272 ± 0.0161 (effect/spread 7.9) and wirings in the 22-burst regime +0.0208 ± 0.0085 (2.4) [exploratory]; each regime separately would have passed the spread clause, and pooling the two regimes is what fails it. The quantity that held across realizations and under intervention is the direction and its modulation, not a universal effect size. Finally, the burst-shortening interventions interact with a separate network-death phenomenon at intermediate parameter values (reported elsewhere); the resulting attrition is tabulated and described in Section 7.6, and is why Section 2.5 reports 4 of 5 wirings and why the milder intrinsic level contributes little to the within-arm dose–response.

## 5. Testable prediction for living cultures

Our original registration made three predictions for dissociated cortical or hippocampal cultures on MEAs (the preparation of Wagenaar et al., 2006; Eytan & Marom, 2006). The first — observational: the stronger half of putative connections shows an excess of pre-before-post spike pairs within population bursts, while first-spike-latency order carries no signal — has now been tested on archived public data and is reported in Section 2.6: the counting half was not reproduced within the stated bounds; the latency half met its negative-only criterion while revealing a positive relationship the model does not predict. The remaining two are interventional, were declared untestable on archived data before that test ran, and are untouched by its outcome in either direction. They are the live prediction, and they are exactly what distinguishes the three readings in the Discussion:

1. An intervention that shortens population bursts at approximately matched burst rate (e.g. modest GABAergic tone increase) should strengthen the strength-ordered pre-before-post excess several-fold, monotonically in realized burst duration. In the model this several-fold amplification is what lifts a sub-resolution excess into a measurable one — so the intervention is also what makes the observational bounded null of Section 2.6 decidable. Two figures should not be confused: the model produces ∼10⁴ pre-before-post pairs per connection in 60 s; resolving an excess of ∼0.2 percentage points at binomial standard error requires of order 10⁵ pairs, so pooling across connections or longer recordings — a statistical requirement of the measurement, not a figure produced by the model. That figure is a binomial floor: pairs share spikes and are not independent, and the empirical burst-shuffle null on the model cohort is about 1.6× wider than the binomial standard error, so the practical requirement is roughly 2–3× larger — the same order of magnitude, stated as a minimum rather than a precision achieved.
2. The effect should track realized burst duration (or per-burst spike count) rather than burst rate, across pharmacologically distinct burst-shortening manipulations. Separating duration from spike count would require a manipulation that moves one without the other, which this model does not contain but a pharmacological preparation might.

What would count as evidence for each reading of Section 2.6: if shortening bursts produces a several-fold, duration-monotone rise in the strength-ordered excess, the effect was present below the observational resolution (reading two); if no rise appears under a strength proxy validated against paired recordings, the mechanism does not transfer (reading one); if the rise appears only once the electrode-level proxy is replaced by a paired-recording estimate of strength, the proxy — not the mechanism — was the limit (reading three).

These require closed-loop or pharmacological intervention on a living culture. Analysis code implementing the instruments of Sections 7.3 and 2.6 is available to collaborating laboratories.

## 6. Preregistration deviations, corrections, and audit trail

Every deviation from a registered protocol, and every correction to a scored number, is listed here. Corrections were found by two independent routes — re-running the registered scorer and independently recomputing from persisted data. Where the two routes agreed, nothing changed; where they disagreed, the disagreement exposed the error (for example, the duplicated rows in the DUR-2 pool). Original values are retained in Supplement S2.

| Issue | When detected | Effect | Treatment |
| --- | --- | --- | --- |
| Tissue-test perimeter (Section 2.6) initially selected six cultures from one plating batch — siblings, not independent preparations by the dataset authors' own conclusion | Before download, during registration | Would have counted one preparation six times | Perimeter amended before download to one culture per batch (six batches), and re-registered |
| DUR-1 scorer voided whole wirings instead of (wiring, level) cells as registered | Before final DUR-1 scoring | Conservative undercount (5/5 read as 3/3) | Corrected to the registered per-cell rule before scoring |
| Tissue-test burst-window function was a sibling of the validated duration function and | After the first scoring pass, before | First scoring pass invalid | Pass withdrawn; entire perimeter re-run; all reported |
| had not itself been validated; it fragmented one burst into ten and collapsed the lag window from ~90 to ~13 ms | results were read as final |  | verdicts are from the corrected run |
| Tissue-test P1 pass rule ($\geq 15/16$) required an effect $\sim 5\times$ the model's own predicted magnitude | After scoring | P1 is a bounded null, not evidence of absence | Reported as a registration error; verdict stands as scored |
| Pairs-per-synapse figure: the per-wiring total ($\sim 3 \times 10^6$) had been reported as the per-synapse count | Before posting, by recomputation from the persisted out-of-sample spikes | Abstract, Results and the schematic overstated the per-synapse count by $\sim 35\times$ | Corrected to the measured value (median 8,960 per synapse; $\sim 3 \times 10^6$ per wiring); the model-produced count and the statistical resolution requirement are now stated separately (Section 5) |
| DUR-2 pooled correlation counted the five shared-baseline cells twice ($n = 23$ instead of 18): the scorer pooled the intrinsic arm, which includes the shared CaS $\times 1.0$ baseline, with all guard-passing DUR-1 cells | After the report, by independent recomputation from persisted data | Pooled $\rho(\text{duration, sorting}) -0.733 \rightarrow -0.790$; $\rho(\text{spikes/burst, sorting}) -0.714 \rightarrow -0.756$; tie-clause gap $0.019 \rightarrow 0.033$ | De-duplicated statistics reported (Section 2.5); originals in Supplement S2; verdict unchanged under either pool; within-arm values unaffected |

## 7. Methods

### 7.1 Neuron and synapse model

Each cell is the single-compartment STG model of Prinz, Billimoria & Marder (2003): eight Hodgkin–Huxley-type currents (I_Na, I_CaT, I_CaS, I_A, I_KCa, I_Kd, I_H, I_leak) with gating kinetics from voltage-clamp fits to lobster STG neurons (Turrigiano et al., 1995; Liu et al., 1998), an intracellular calcium pool (τ_Ca = 200 ms), and the calcium reversal potential recomputed each step from the Nernst equation with [Ca]_ext = 3 mM at T = 283 K (10 °C, the standard STG bath temperature). Excitatory cells use a validated burster conductance set (AB/PD-type), inhibitory cells a validated LP-type set; both sets were taken from a separately validated pyloric-circuit implementation rather than hand-tuned for this study. Baseline synapses between identified pyloric neurons are graded (Prinz, Bucher & Marder, 2004); the plastic recurrent connections described below are spike-mediated.

### 7.2 The simulated culture

The “dish” comprises 48 excitatory and 12 inhibitory neurons, modelled as a small conductance-based network in the STG tradition (Abbott & Marder, 1998). Plastic excitatory→excitatory connections are drawn randomly at connection probability 0.15, with the total incoming conductance budget fixed at (N−1)·p·w̄ (w̄ = 0.015) so that wirings differ in structure, not in total drive. Inhibitory→excitatory conductance g_ie is a global parameter. Short-term depression (release fraction 0.20, recovery τ = 400 ms) sets whether the network bursts at all rather than latches into sustained firing: without it this cell model does not produce culture-like population bursts. (What terminates an individual burst was not established here; Section 2.5 shows an intracellular conductance controls burst duration.) Plasticity is pair-based additive STDP (no weight dependence; hard bounds w ∈ [0, 0.06]) with exponential kernels, A₊ = 0.012, τ₊ = 16.8 ms, τ₋ = 33.7 ms (the time constants of Bi & Poo, 1998), and A₋ solved from the area-ratio constraint below; additive STDP with hard bounds is the unstable end of the STDP-model spectrum (van Rossum, Bi & Turrigiano, 2000; Morrison, Diesmann & Gerstner, 2008), which makes the neutral-point construction below load-bearing. Initial weights are drawn at the budget mean with 20% Gaussian jitter, clipped at zero.

We defined the neutral point from the integral of the STDP window rather than from zero net change (integral −0.134, Song, Miller & Abbott, 2000; A₋/A₊ = 0.8299 in closed form). This modelling choice was fixed before the experiments, held constant across all conditions, and not fitted to the present results; its sensitivity was not tested. Zero-area perturbations around the rule’s own ratio, rather than around zero, serve as the null. The provenance of the constant and the registered experiment that motivated the construction are described in Supplement S1.

Integration is exponential Euler (Dayan & Abbott, 2001) at dt = 0.05 ms. Spontaneous activity is endogenous — the excitatory conductance set is an intrinsic burster and no noise or external drive is injected. Each run simulates 60 s of biological time. A network “wiring” is the random structure generated by one seed; wirings are the unit of analysis throughout, as independent connectivity realizations of one model. Spike times, weights, class labels and burst windows are persisted per run.

#### Computational resources

Runs cost ≈3.7 s wall-clock per biological second on a single CPU core; every experiment reported here was run single-threaded on a consumer-grade CPU.

#### Software

The simulator, plasticity rule, and burst-detection code are custom NumPy implementations (exponential Euler, dt = 0.05 ms); no external simulator was used. No validation against Brian2 or NEURON was performed: the single-cell model was validated only against the bursting dynamics reported by Prinz et al. (2003), and the burst-duration instrument of Section 2.6 against the published Wagenaar et al. (2006) values.

### 7.3 Instruments and definitions

Bursts are detected in 10 ms bins at a 20% active-fraction threshold, merged across gaps below 50 ms, then delimited at full width at 20% of each burst’s own peak at 1 ms resolution (Figure 1A).

Synapses are split at the median conductance per wiring into equal strong and weak classes (Figure 1B). For each class we compute P/M, the ratio of kernel-weighted potentiation to depression pressure over pairs with |lag| ≤ 250 ms inside burst windows, read against the neutral point rather than zero; and the pre-before-post pair fraction, the proportion of pre-before-post spike pairs among all pairs — a pure count with no kernel weighting. “Pre-before-post” denotes the temporal order that a pair-based STDP rule potentiates; the term asserts nothing about whether the presynaptic spike caused the postsynaptic one. The outcome measure is the strong-minus-weak difference in each quantity.

We avoid the term *consolidation*. What is measured is an increase in the separation of the pre-existing synaptic weight distribution between stronger and weaker connections under spontaneous activity, over the 60 s simulated; we call this *short-timescale synaptic weight sorting*, or simply *sorting*, and do not imply long-term stabilization or memory consolidation in vivo.

### 7.4 Preregistration

Predictions, decision trees, priors, and falsifiers were written into a version-controlled record and committed before the corresponding simulations ran (the out-of-sample replication was pushed to the repository before the first simulation step). Criteria are scored as written; where a criterion’s clauses split, the split is reported. Post-hoc observations are labeled as such and used only to seed the next registration, never scored retroactively. Instruments are validated against published values of the target preparation before scoring, and the validation must cover the code that actually runs. Deviations and corrections are listed in Section 6. Numbers that were not scored under a registration — post-hoc splits, descriptive correlations, calibration checks — are marked [exploratory] where they appear. The full registration record is summarized in Supplement S2.

### 7.5 Statistics

Wirings (or recordings) are the unit of analysis; wirings are independent connectivity realizations of one model, and replication is reserved for the out-of-sample cohort. Directional predictions are scored by the count of wirings in the predicted direction against the pre-specified bar, with an exact binomial test reported where the bar is all-or-nearly-all (for 10 of 10 under a fair coin: one-sided p = 2⁻¹⁰ = 0.000977, two-sided 0.00195). The test assumes independence between wirings; connectivity draws of one model system share every other parameter, so it is a test of direction consistency, not of independent preparations. Means with t-statistics are reported as secondary descriptions of magnitude. Correlations are Spearman’s ρ unless stated.

### 7.6 Exclusions and voids

Every exclusion follows a rule fixed in the corresponding registration before running: a dead network or a failed guard voids that wiring × level cell only, never the wiring. The complete attrition is tabulated here so that no result is read without it.

| Experiment | Unit | Planned | Valid | Voided | Reason for voiding |
| --- | --- | --- | --- | --- | --- |
| stg-P/M<br>(Sections 2.1–2.2) | wiring | 10 | 10 | 0 | None. |
| PM-4 (Section 2.3) | wiring | 10 | 10 | 0 | None. |
| DUR-1 (Section 2.4) | (wiring, g_ie level) | 15 | 13 | 2 | Registered rhythm guard: fewer than 3 bursts, or one burst spanning >20% of the analysis window. Wirings 30 and 34 at g_ie = 0.06 only. The scored comparison (0.03 vs 0.10) does not use that level, so all 5 wirings entered it. |
| DUR-2 (Section 2.5) | (wiring, g_CaS level) | 10 | 5 | 5 | Same rhythm guard. Voided: wiring 30 at 0.8; 32 at 0.8; 33 at 0.6 and 0.8; 34 at 0.8. The g_CaS = 1.0 condition is DUR-1's g_ie = 0.03 baseline and was not re-run. |
| BAND-1 | (wiring, level), two levers | 56 | 56 | 0 | None. 15 of 56 cells reached the silent absorbing state (7 on the g_ie lever, 8 on g_CaS); that is the measured outcome, not an exclusion. |
| MEA-1 (Section 2.6) | recording | 18 | 16 | 2 | Registered guard of at least 8 detected bursts: 2-1-10 (1 burst) and 2-1-17 (4 bursts). |

In DUR-1 the void is per cell (wiring, level), not per wiring (Section 6 records the scorer correction). In DUR-1 the rate guard (burst-rate spread < 10%) was met (8.6%) and voided nothing.

What the rhythm guard detects is a network-death phenomenon: a network fires normally for some seconds and then falls permanently silent, with synaptic weights intact. It is detected by the burst-count guard above and handled by the per-cell void rule. It removes more wirings at *intermediate* parameter values than at the extremes — in the intrinsic arm four of five at the milder level (×0.8) against one of five at the stronger (×0.6) — a non-monotone attrition characterized separately (BAND-1, reported elsewhere). The void rule cannot favor the hypothesis: a voided cell is absent from both sides of every scored contrast, and the scored DUR-1 contrast never touches the voided level.

## Author contributions

F.G. designed and ran the study, performed the analyses, and wrote the manuscript. J.F. contributed to the conception and interpretation of the work through continuous discussion and revised the manuscript. Both authors approved the final version.

## AI assistance statement

Generative AI tools (Claude, Anthropic) were used under the authors’ direction for portions of code development, data-analysis support, figure-generation workflows, and manuscript drafting. The authors independently reviewed and verified the analyses, figures, claims, references, and final text and retain full responsibility for the work. No AI system is an author of this manuscript.

## Competing interests

The authors are founders of Luviner, a company developing edge machine-learning products. The work reported here concerns a separate biophysical simulation line; no Luviner product is evaluated or promoted in this manuscript.

## Funding

This work received no external funding and was supported internally by Luviner. J.F. holds a research fellowship at the University of Parma unrelated to this work.

## Data and code availability

Code, registration records, per-run results, the MEA analysis pipeline with a checksummed data manifest, and figure-generation scripts are publicly available at https://github.com/luviner-ai/luviner-neurolab (MIT license for code, CC-BY 4.0 for text and figures) and archived at Zenodo, DOI 10.5281/zenodo.22287793 (release v1.0.1). The curated laboratory record (`LAB-RECORD.md`) lists, for each registered experiment, the hash of the commit in the private working record that introduced it; that record is available to reviewers on request. The tissue test of Section 2.6 uses the publicly available Wagenaar, Pine & Potter (2006) dataset, which is not redistributed; the manifest carries its URLs and checksums, and its registration was committed before download.

## Supplementary Methods S1

### S1.1 The neutral point

The STDP window integral was fixed at −0.134 — the standard net-depressing choice (Song, Miller & Abbott, 2000) — and A₋ follows in closed form from A₊, τ₊ and τ₋, giving A₋/A₊ = 0.8299. The value is not a measurement: no data enters it, and it was held fixed across all conditions and experiments, so the reference against which sorting is read is the same constant throughout (verified by reading the rule object under the extreme conditions of each experiment). What a separate registered experiment did establish — on a different, cortical-style preparation, where it measured a neutral ratio of 0.605 ± 0.046 against a predicted 0.598 — is the principle that the dividing line for sorting is the rule’s own area ratio rather than zero net area; that principle, not that value, is what is applied here. The constant happens to lie inside the range of this preparation’s own P/M ratios (0.776–0.888), an observation made afterwards, not the way it was chosen. The construction is BCM-adjacent (Bienenstock, Cooper & Munro, 1982) in that a fixed potentiation/depression balance sets a threshold that the preparation’s activity is read against. The sensitivity of the qualitative findings to the fixed integral has not been measured (Section 4).

### S1.2 The burst-shuffle null and its asymmetry

The per-pair null for Section 2.6 pairs unit A’s spikes from burst *i* with unit B’s from burst *j* (i ≠ j), over 100 derangements. It preserves spike count, firing rate, and each unit’s typical position and shape within a burst — every property that could shift a baseline off 0.5 — and destroys only same-burst coordination. It deliberately *retains* any static lead–lag between two units: if A tends to fire before B in every burst, the shuffle keeps that, because Section 2.1 found no support for static order as the associated statistic, and a null that removed it would credit the effect for the very thing not shown to carry it. The consequence is an asymmetry stated before the numbers: the null subtracts any synapse-induced static phase shift, so the measured excess is a lower bound on same-burst coordination — it can miss true signal but cannot manufacture it. A spike-time jitter null was considered and rejected before download: at the scale of the lag window (half a burst), jitter wide enough to matter makes both trains uniform and collapses the null to the nominal 0.5 for every pair regardless of rate, while jitter narrow enough to preserve the lag distribution leaves the null on top of the data — a fine-timing instrument applied to a coarse-ordering statistic.

## Supplement S2 — Registration record

Each row summarizes what was committed before the corresponding run, and what was scored. Commit hashes ([hash]) will be filled from the public repository at posting.

| Experiment | Prediction | Criterion | Falsifier | Outcome | Deviations / voids |
| --- | --- | --- | --- | --- | --- |
| stg-P/M<br>(Sections 2.1–2.2) [hash] | Mechanism:<br>strongly driven cells fire earlier.<br>Outcome: bursts favor strong synapses. | Mechanism:<br>$\rho$(latency, incoming weight) negative in $\geq 8/10$ wirings.<br>Outcome:<br>strong–weak P/M positive in $\geq 8/10$ **and** effect > between-wiring spread. | Mechanism criterion unmet → temporal hypothesis not supported.<br>Outcome sign clause unmet → sorting absent. | Mechanism 5/10 → not met.<br>Outcome sign 10/10 met; spread 0.92 → not met. | None; 0 voids.<br>Post-hoc regime split noted, not scored. |
| PM-4<br>(Section 2.3) [hash] | Direction replicates on 10 wirings that did not yet exist. | (1) P/M difference positive $\geq 9/10$; (2) pre-before-post excess positive $\geq 9/10$; (3) $\rho$ between measures $\geq 0.60$. | Any clause unmet → direction not replicated. | 10/10, 10/10, $\rho = +0.988$ → all met (one confirmation, not three: ratio CV 0.13). | Registration pushed before the first simulation step; 0 voids. |
| DUR-1<br>(Section 2.4)<br>[hash] | Shorter bursts sort more strongly (direction fixed in advance). | P1: pre-before-post excess larger at g_ie 0.10 than 0.03 per wiring; P2: P/M difference larger; guard: rate spread <10% (failure voids the cell). | P1/P2 pass while the observational arm P3 fails → evidence for g_ie, not duration. | P1 5/5, P2 5/5; rate spread 8.6%. P3 uninformative (444–449 ms, 1.0% spread) — falsifier did not fire. | Two cells voided at g_ie 0.06 (network death). Scorer initially voided whole wirings, corrected to per-cell as registered (Section 6). |
| DUR-2<br>(Section 2.5)<br>[hash] | Intrinsic shortening (g_CaS) strengthens sorting; sorting tracks realized duration pooled and within each arm. | P1: stronger at $\times 0.6$ than $\times 1.0$ within wirings; P2: $\rho(\text{duration, sorting})$ pooled and within each arm; P3: within intrinsic arm, shortest beats longest. Ceiling: mediator is "duration, or the spike count that falls with it"; tie-clause fires if $\rho(\text{spikes/burst})$ matches or beats $\rho(\text{duration})$. | Intrinsic shortening does not strengthen sorting → "duration" dies, result belongs to inhibition. | P1 4/4 (4 of 5); P2 –0.790 (de-duplicated, n=18; the scorer as registered printed –0.733 on a pool that counted the shared baseline twice) / –0.648 / –0.808; P3 met. Tie-clause missed by 0.033 (0.019 as originally scored) → ceiling stands verbatim. | $\times 0.8$ voided 4/5, $\times 0.6$ voided 1/5 (network death); one wiring lost both levels. Baseline shared with DUR-1 by design; pooled statistic de-duplicated after the report (Section 6). |
| BAND-1<br>[hash] | Cross-reference: the network-death phenomenon at intermediate parameter values, reported elsewhere. | — | — | — | Appears here only as attrition (Section 7.6). |
| MEA-1<br>(Section 2.6)<br>[hash] | Observational half of Section 5 on Wagenaar 2006: strong half shows higher pre-before-post fraction; sorting tracks duration negatively within age strata; latency carries no negative signal. | P1: $\geq N-1$ of N recordings; P2: negative within-stratum correlation, with an underpower clause (within-stratum spread < 25% of across-stratum → underpowered, not refuted) and a unit-count tie-clause (→ unresolved); P3: no consistently negative | P1 unmet → the counting mechanism does not appear under an independent strength proxy. P2 unmet with neither clause firing → duration relation absent on the merits. | P1 6/16 → not met (bounded null: CI [–0.53, +0.32] pp). P2 +0.08, clauses did not fire → not met. P3 met (descriptive positive relation, unscored). | Perimeter amended before download from batch siblings to one culture per batch (6 batches). First scoring pass withdrawn (unvalidated burst-window function) and perimeter re-run. 2 voids (<8 bursts). P1 bar |
|  |  | latency relation.<br>Guard: <8 bursts<br>voids the recording. |  |  | later found<br>unpassable at the<br>model's own<br>effect size —<br>registration error,<br>reported (Section<br>6). |

## Supplementary Results S3 — Exploratory checks and the DUR-2 tie-clause

### S3.1 Post-hoc checks on the out-of-sample cohort

Two checks were added after the results were seen; neither was in the PM-4 registration, and neither can be read at the registered bar. Both recompute from the spikes, burst windows, wiring and weights persisted by the out-of-sample runs, with no new simulation; the analysis module first reproduces the persisted per-wiring sorting and pre-before-post excesses from those raw fields, with zero discrepancy, before computing anything new. The original cohort did not persist synapse endpoints, so both checks are restricted to the ten out-of-sample wirings.

*Weight as a continuous predictor* (Figure S1). PM-4 registered a median split into strong and weak halves. Replacing the split with the graded weight, the rank correlation between a synapse’s weight and its pre-before-post pair fraction is ρ = +0.386 averaged across wirings, positive in 10 of 10 [exploratory]. The median across-wiring curve of pair fraction against weight decile rises with weight but is not monotone [exploratory]; two wirings (seeds 17 and 18) sit far from the modal bursting regime and carry gaps five to ten times the rest, which is why the summary is the median rather than the mean. A 25/75 quantile contrast, chosen after the fact, gives +0.505 percentage points against the registered median split’s +0.346, positive in 10 of 10 for both [exploratory] — a sharper contrast produces a larger gap, as a graded relationship would predict.

*Kernel ablation* (Figure S2). The quantity reported in Section 2.3 is the coefficient of variation, across wirings, of the ratio between the kernel-weighted sorting measure (P/M) and the flat pre-before-post count excess; it is reproduced here at 0.132 against the recorded 0.13. A different quantity, the per-synapse rank correlation between count fraction and kernel-weighted P/M, is ρ = +0.977 [exploratory]; it is reported as an additional check with its definition stated and is not the across-wiring CV. Both say the same thing at different grain: under the present STDP rule and activity regime, the exponential kernel reorders almost nothing relative to the bare count. The departure from proportionality is visible but not established: ρ(cf_diff, ratio) = −0.358, permutation p = 0.31 (20,000 shuffles); seed 17 carries the largest standardized residual (z = −2.74) and most of the CV (0.132 → 0.093 without it) [exploratory].

*Per-synapse null* [exploratory, post-hoc]. The burst-shuffle null that the tissue test registered before its download (Section 2.6; Supplement S1) was applied per synapse to the PM-4 cohort, 50 derangements, no simulation. The surrogate is unbiased: its own strong-minus-weak gap has median −0.0001 percentage points, positive in 5 of 10 wirings, so the raw excess is not an artifact of the null’s construction. The excess *over* the null has median +0.022 percentage points, positive in 8 of 10 wirings, two-sided sign test p = 0.109 [exploratory] — suggestive, not established at n = 10.

The mean (+0.383 points) is carried entirely by the two off-regime wirings (seeds 17 and 18); the median is the summary that survives them. This per-synapse statistic is a different estimator from the registered pooled count excess, and its 8 of 10 does not contradict the registered 10 of 10.

Holding Sections 2.1–2.5 to the standard of Section 2.6 weakens them, from 10 of 10 at the registered estimator to 8 of 10 at p = 0.11 under a per-synapse surrogate; this is reported rather than omitted.

### S3.2 The DUR-2 tie-clause and its bootstrap

#### The pre-specified ceiling stands

Spikes per burst falls with duration in both arms (e.g. 1273 → 385 in one wiring), and the tie-clause — fire if ρ(spikes/burst, sorting) matches or beats ρ(duration, sorting) — missed by 0.033 (−0.756 vs −0.790) on 18 de-duplicated points (0.019 as originally scored on the duplicated pool), which is within noise at this n. A bootstrap over the 18 cells (10,000 resamples; the two correlations are dependent, sharing rows and with predictors correlated at ρ = +0.93) puts |ρ(spikes/burst)| − |ρ(duration)| at −0.033 with 95% CI [−0.186, +0.089] [exploratory]: the two candidate mediators are indistinguishable at this n, and in 27.4% of resamples the registered tie-clause would have fired and declared the result unresolved. Duration was therefore not ruled out as the mediator, but neither was it established; the gate passed narrowly, and we report the margin beside the verdict. (The registered clause compares magnitudes; only that convention is reported here.) The claim the registration licenses is therefore, verbatim: the mediator is **“burst duration, or the spike count that falls with it.”** Separating duration from spike count would require a lever that moves one without the other, absent in this model.

---

**Figure S1.**
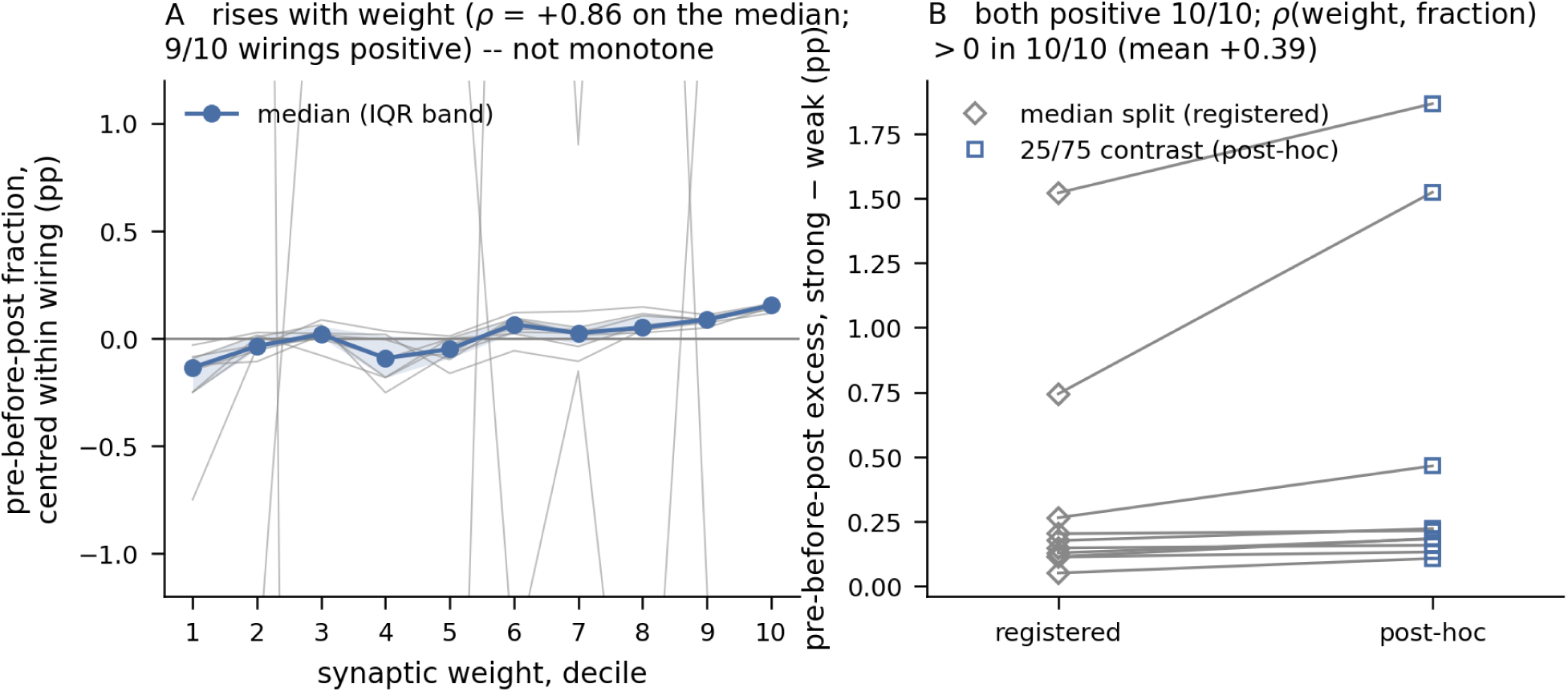
Weight as a continuous predictor [exploratory, post-hoc — not registered]. (`figS1_continuous_weight.pdf`, from the persisted out-of-sample runs via `experiments/dish/pm4_posthoc.py`.) **A**, pre-before-post pair fraction, centred within each wiring, against synaptic weight decile (deciles by rank, since many weights are identical at the bound and near zero). Thin lines, individual wirings; the summary is the **median** with the interquartile band, not the mean: two wirings (seeds 17, 18) sit far off the modal regime with gaps five to ten times the rest, and the mean curve is not monotone even in the direction of its own rank correlation. The median rises with weight and is not monotone either. **B**, paired on one axis: the registered median-split excess (open diamonds) and the post-hoc 25/75 contrast (filled) per wiring; both positive in 10 of 10.

**Figure S2.**
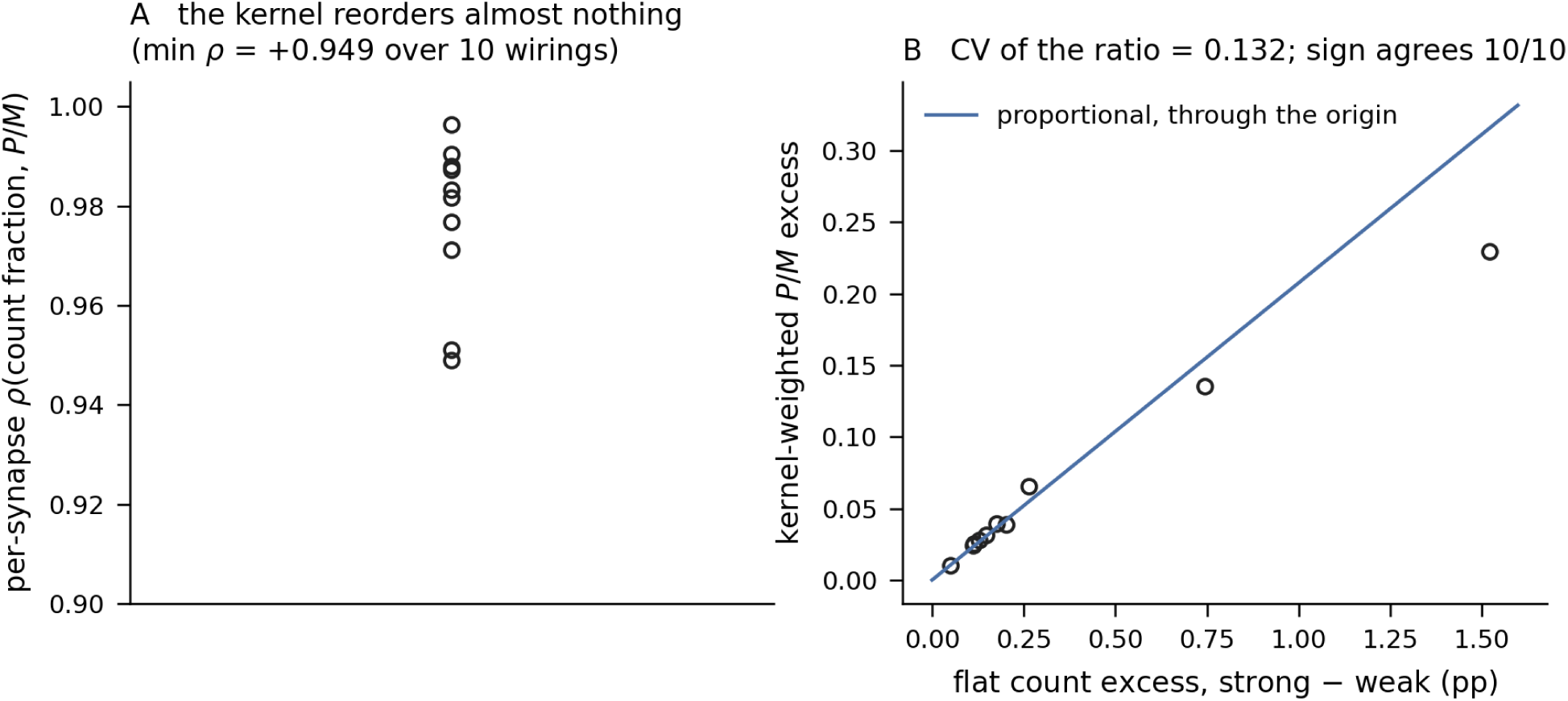
Kernel ablation: flat count versus exponential kernel [exploratory, post-hoc — not registered]. (`figS2_kernel_ablation.pdf`, same source.) **A**, per-synapse rank correlation between the flat pre-before-post count fraction and the kernel-weighted P/M, one point per wiring (mean +0.977). **B**, kernel-weighted P/M excess against flat pre-before-post count excess (strong − weak), one point per wiring, with the proportional line through the origin at the mean per-wiring ratio; the across-wiring CV of that ratio is 0.132 (Section 2.3 reports 0.13), and the sign agrees in 10 of 10

## Notes

https://github.com/luviner-ai/luviner-neurolab

https://doi.org/10.5281/zenodo.22287793

### Competing Interest Statement

The authors are founders of Luviner, a company developing edge AI machine-learning products. The work reported here concerns a separate biophysical simulation line; no Luviner product is evaluated or promoted in this manuscript.

